# Extra-cellular matrix induced by steroids through a G-protein coupled receptor in a Drosophila model of renal fibrosis

**DOI:** 10.1101/653329

**Authors:** Wenjing Zheng, Karen Ocorr, Marc Tatar

## Abstract

Aldosterone is produced by the mammalian adrenal cortex to modulate blood pressure and fluid balance, however excessive, prolonged aldosterone production promotes fibrosis and kidney failure. How aldosterone triggers disease may involve actions that are independent of its canonical mineralocorticoid receptor. Here we present a *Drosophila* model of renal pathology caused by excess extra-cellular matrix formation, stimulated by exogenous aldosterone and insect ecdysone steroids. Chronic administration of aldosterone or ecdysone induces expression and accumulation of collagen-like pericardin at adult nephrocytes – podocyte-like cells that filter circulating hemolymph. Excess pericardin deposition disrupts nephrocyte (glomerular) filtration and causes proteinuria in Drosophila, hallmarks of mammalian kidney failure. Steroid-induced pericardin arises from cardiomyocytes associated with nephrocytes, reflecting an analogous role of mammalian myofibroblasts in fibrotic disease. Remarkably, the canonical ecdysteroid nuclear hormone receptor, ecdysone receptor EcR, is not required for aldosterone or ecdysone to stimulate pericardin production or associated renal pathology. Instead, these hormones require a cardiomyocyte-associated G-protein coupled receptor, dopamine-EcR (dopEcR), a membrane-associated receptor previously characterized in the fly brain as affecting behavior. This *Drosophila* renal disease model reveals a novel signaling pathway through which steroids may potentially modulate human fibrosis through proposed orthologs of dopEcR.

**Significance Statement:** Aldosterone regulates salt and fluid homeostasis, yet excess aldosterone contributes to renal fibrosis. Aldosterone acts through a nuclear hormone receptor, but an elusive, G-protein coupled receptor (GPCR) is thought to also mediate the hormone’s pathology. Here we introduce a Drosophila model of renal fibrosis. Flies treated with human aldosterone produce excess extra-cellular matrix and that causes kidney pathology. Flies treated with the insect steroid ecdysone produce similar pathology, and from this analogous response we identify an alternative receptor through which steroids mediate renal fibrosis -- the GPCR dopamine-Ecdysone Receptor (dopEcR). dopEcR functions in heart muscle cells associated with nephrocytes, analogous to the role of myofibroblasts in human fibrosis. This finding opens avenues to identify mammalian GPCR homologs of dopEcR through which aldosterone mediates renal fibrosis.

## Introduction

Aldosterone is a primary regulator of sodium and potassium homeostasis in kidney distal tubules. But when chronically elevated as in diabetes and primary aldosteronism (1), aldosterone promotes kidney interstitial fibrosis and glomerulosclerosis (2–4). These events are preceded by elevated inflammation through monocytes and macrophage infiltration followed by proliferation of myofibroblasts that secrete fibrinogen, collagens and elastins. Aldosterone increases reactive oxygen species (ROS) to induce profibrotic factors such as Transforming Growth Factor-β1 (TGF-β1), Plasminogen Activator Inhibitor-1, and Enothelin-1 (4). TGF-β1 contributes to fibrosis by activating myofibroblasts (5) as well as through suppressing matrix metalloproteinases, which can further promote excess extra-cellular matrix (6). Aldosterone affects these processes through its interaction with the mineralocorticoid nuclear hormone receptor (MR), as inferred from studies where blockade of MR activity prevents aldosterone-associated inflammatory and fibrotic outcomes (7–9).

Many data also suggest that aldosterone contributes to fibrosis through rapid signaling independent of MR (4). Aldosterone enhances TGF-β1 expression and fibrosis in part through stimulation of ERK1/2 that cannot be blocked by spironolactone, (10–12), while aldosterone fosters hypertrophy in cardiomyocytes through action on ERK5 and PKC (13). As well, aldosterone effectively induces calcium influx in fibroblasts that are derived from MR-deficient mice (14). Angiotensin receptors crosstalk with MR to modulate NF-κB in vascular smooth muscle cells (VSMC) stimulated with aldosterone (15), suggesting that aldosterone may act through G-protein-coupled receptors (GPCR). With considerable debate, GPER1 has been proposed as an alternative GPCR for aldosterone (16–19). In VSMC, aldosterone was seen to activate PI3 kinase and ERK through both GPER1 and MR (20). Emerging evidence, however, shows that 17β-estradiol is the steroid agonist of GPER1 (21–23), and no pharmacological evidence demonstrates GPER1 to interact with aldosterone. The problem remains: through which receptor aside from MR might aldosterone stimulate G-coupled signaling and how does this modulate fibrosis?

Here we address these issues with a new model of steroid-induced fibrosis based on *Drosophila melanogaster*. Genetic data reveal the GPRC dopamine-EcR (dopEcR) is necessary and sufficient for aldosterone and insect ecdysone to induce excess extra cellular matrix at nephrocytes and to disrupt fly renal function. Based on our findings we propose that mammalian homologs of dopEcR may offer a novel entrée to moderate fibrotic pathology in humans.

## Results

The tubular heart of adult *Drosophila* is surrounded by a collection of pericardial cells, podocyte-like nephrocytes that conduct size-selective filtration of hemolymph (24, 25). The heart tube and the associated nephrocytes are enmeshed in an extracellular matrix composed of collagen-like proteins including pericardin (collagen IV) (26, 27). In a first step to develop a model of *Drosophila* renal fibrosis we quantified nephrocyte function my measuring levels of protein in excreta (frass) as an analog to proteinuria seen in humans with glomerular dysfunction (28). Frass arises as a by-product of digestion and from discharge of Malpighian renal tubules. Frass quality is modulated in response to diet, mating and internal metabolic state (29), and in response to activity of nephrocytes (30, 31). We collected frass from adult males in microcentrifuge tubes and assayed for total protein content normalized to uric acid as a measure of excretion volume. Diets of high sugar or salt decreased protein excretion compared to normal diet (Fig 1A). Protein in frass was elevated in adults fed aldosterone for two weeks (Fig 1B) yet not when fed aldosterone for only 24 h (Fig S1). To determine if nephrocytes contribute to frass protein content, we depleted nephrocyte slit diaphragm genes *kirre* and *sticks-n-stones* (*sns*), which encode homologs of mammalian nefrin. Previous reports show that reduced *kirre* and *sns* impairs nephrocyte filtration measured by uptake of fluoro-dextran beads (25, 32). We replicate this result (Fig 1D, E) and subsequently observed that reduced *kirre* and *sns* also elevates protein excretion (Fig 1C). Thus, defects in nephrocyte function are sufficient to induce proteinuria in *Drosophila*.

**Figure 1.**
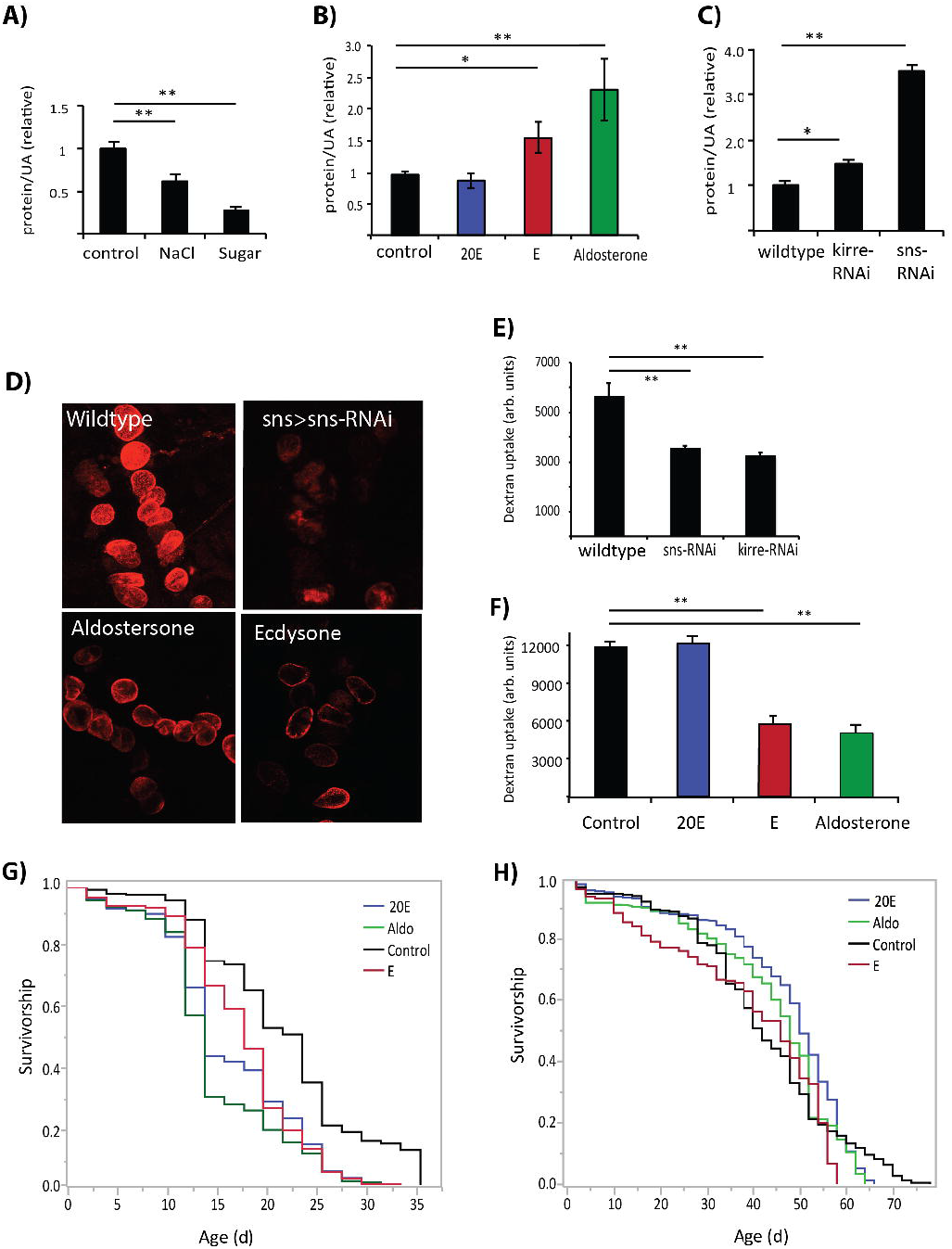
Aldosterone and ecdysone induce renal dysfunction in adult Drosophila. **A)** Proteinuria in 3 w old males measured as excreted protein/uric acid in adults fed high salt diet and high sugar diet, assessed across four independent wildtype backgrounds, each with 4-5 biological replicates. Values normalized to control treatment within each background. **B)** Proteinuria in 3 w old males fed for two weeks with 20-hydroxyecdysone (20E), ecdysone (E) or aldosterone, assessed with three wildtype backgrounds, each with 4-5 biological replicates. **C)** Proteinuria in 3 w old males expressing RNAi in nephrocytes to deplete slit diaphragm encoding *kirre* or *sns*. **D)** Confocal images (representative z-stack) of nephrocytes of 3 w old females in *ex vivo* dextran-bead filtration assay. Efficient filtration presented by wildtype; impaired filtration occurs with depletion of slit diaphragm (sns-RNAi) and by treatment of wildtype with aldosterone or ecdysone. **E, F)** Quantification of fluorescence intensity from biological replicates of nephrocytes in *ex vivo* dextran-bead filtration assay when slit diaphragm is depleted by RNAi, and wildtype adults are treated with 20E, ecdysone or aldosterone. (A-C, E, F: ANOVA with Dunnett’s post hoc comparison to control, * p < 0.05, ** p < 0.001. Plots show means with SD.) G) Survival upon instant diet supplemented with NaCl (1.5%) for cohorts (each N= 230-330) while treated with 20E, ecdysone, or aldosterone relative to control. Survival was reduced in each treatment, pair-wise contrasts to control, log-rank test, p < 0.001. **H)** Survival upon instant diet for cohorts (each N= 216-280) while treated with 20E, ecdysone, or aldosterone. Relative to control (median life span = 42 d), survival was increased by 20E (median life span = 50 d; log-rank test, p = 0.051), but not significantly affected by aldosterone (median lifespan = 48 d, log-rank test, p = 0.742) or ecdysone (median lifespan = 46 d, log-rank test, p = 0.185).

*Drosophila* do not synthesize aldosterone, a mammalian steroid hormone produced in the renal cortex. Aldosterone may act in *Drosophila* either as a mimic of insect steroids or by providing a precursor in the synthesis of insect steroids. Notably, 20-hydroxyecdyone (20E) is the primary active steroid in *Drosophila*. 20E is oxidized from prohormone ecdysone by 20-hydroxylase (encoded by *shade)* at target cells. 20E activates the nuclear hormone ecdysone receptor (EcR) to modulate transcription. Feeding adults 20E for two weeks did not stimulate proteinuria, but proteinuria was elevated in adults chronically fed ecdysone (Fig 1B). Likewise, chronic aldosterone and ecdysone, but not 20E, suppressed dextran filtration by nephrocytes (Fig 1F). While aldosterone and ecdysone affect nephrocyte function and associated proteinuria, all tested steroids (aldosterone, ecdysone and 20E) reduced the survival of adults on high salt diet (Fig 1G). We found no consistent association between exogenous steroids, renal function and survival of adults on normal diet (Fig 1H).

The extracellular matrix surrounding pericardial nephrocytes consists of collagen-like proteins including pericardin, col4a1 and Viking (26, 27, 33). Pericardin (*prc*) mRNA was induced by overnight feeding with aldosterone and ecdysone, but not by 20E (Fig 2A). Neither *col4a1* nor *Viking* mRNA were induced by any of the tested steroids (Fig 2B, C). Despite induction of *prc* mRNA, overnight steroid feeding itself did not elevate proteinuria (Fig S1). In contrast, aldosterone and ecdysone fed to wildtype adults for two weeks stimulated elevated pericardin protein deposition in the nephrocyte extracellular matrix, while no effect was seen with 20E (Fig 2D, E). Depletion of *pericardin* mRNA from cardiomyocytes *(tinΔ4-* gal4>prc(RNAi)) but not nephrocytes (sns-gal4>prc(RNAi)) blocked the ability of aldosterone and ecdysone to induce excess pericardin deposition (Fig 2D, E). Furthermore, pericardin stimulated by aldosterone and ecdysone is sufficient to produce proteinuria and impair nephrocyte filtration. Depletion of *prc* mRNA from cardiomyocytes blocks the ability of aldosterone and ecdysone to induce pathology, while driving prc(RNAi) in nephrocytes does not (Fig 2F-K). Thus, cardiomyocytes are the source of pericardin protein that accumulates in response to chronic exposure to aldosterone and ecdysone, and impairs nephrocyte function.

**Figure 2.**
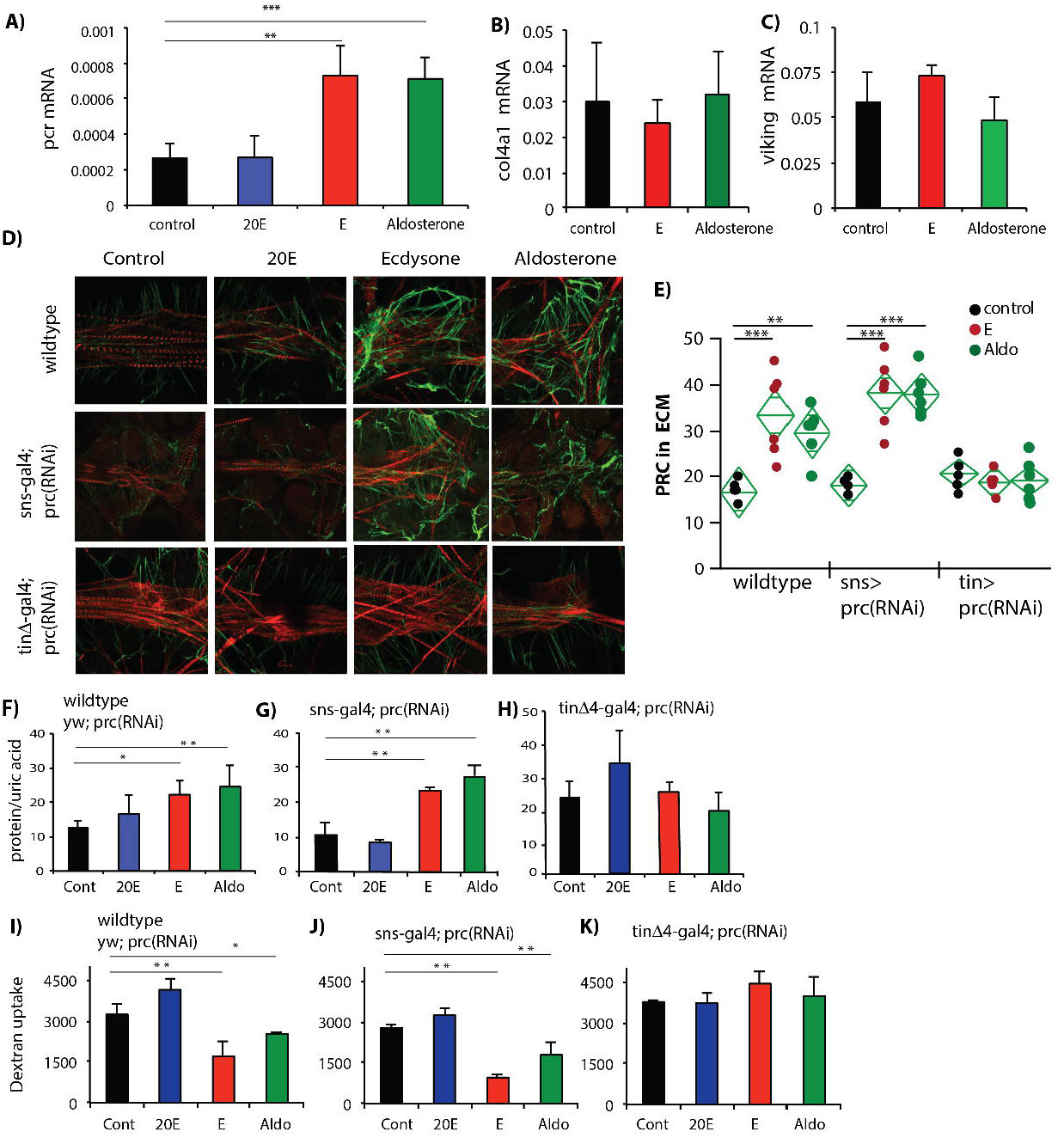
Pericardin from cardiomyocytes induced by steroids produces renal dysfunction. **A)** *Pericardin* (prc) mRNA in nephrocyte-cardiac tissue induced by ecdysone (E) and aldosterone, but not by 20-hydroxyecdysone (20E). **B, C)** *Collagen-4a1 (col4a1)* and *Viking* mRNA in nephrocyte-cardiac tissue are not induced by steroid hormones. **D)** Confocal images (representative z-stacks) of nephrocyte-cardiac tissue of 3 w old females after two-week treatment with 20-hydroxyecdysone, ecdysone or aldosterone; wildtype and knock-down genotypes to deplete *prc* mRNA in nephrocytes (*sns*-gal4>UAS-*prc*(RNAi)) and cardiomyocyte (*tin*Δ4-gal4>UAS-*prc*(RNAi)). Phalloiden (red) stains cardiomyocyte actin; secondary antibody labels (green) pericardin protein in extra-cellular matrix about nephrocytes and heart. **E)** Quantification of straining intensity for pericardin ECM protein (each with six independent nephrocyte-heart preparations). **F-H)** Proteinuria in 3 w old males fed for two weeks with 20-hydroxyecdysone, ecdysone or aldosterone, assessed in wildtype background (yw/UAS-*prc*(RNAi)), and in genotypes that reduce *pericardin* (UAS-*prc*(RNAi)) in nephrocytes (*sns*-gal4) or cardiomyocytes (*tin*Δ4-gal4). **I-K)** Quantification of fluorescence intensity from biological replicates of nephrocytes in *ex vivo* dextran-bead filtration assay in 3 w old males fed for two weeks with 20-hydroxyecdysone, ecdysone or aldosterone, assessed in wildtype (yw/UAS-*prc*(RNAi), and in genotypes that reduce *pericardin* (UAS-*prc*(RNAi)) in nephrocytes (*sns*-gal4) or cardiomyocytes (*tin*Δ4-gal4). (A-C, E-K: One-way ANOVA with Dunnett’s comparison relative to control, * p < 0.05, p < 0.01. Plots show mean with SD)

It is striking that ecdysone but not 20E induces pericardin and associated renal pathology in adult *Drosophila*. This suggests that *prc* expression and its accumulation in the ECM can be regulated independent of EcR, the canonical nuclear hormone ecdysone receptor of 20E. Indeed, depletion of *EcR* by RNAi in cardiomyocytes did not prevent the steroid-dependent induction of *prc* mRNA, or associated ECM accumulation and renal pathology (Fig 3A, C, E). As an alternative, dopamine-EcR (dopEcR) is a membrane GPCR receptor of ecdysone described to function in the fly brain (34, 35). We found that *dopEcR* mRNA could be detected in adult pericardial tissue (heart-nephrocytes), and these levels increased after feeding with aldosterone and ecdysone (Fig 3G). Consistent with the notion that dopEcR is required for aldosterone and ecdysone to stimulate renal pathology, depleting *dopEcR* from cardiomyocytes blocks the ability of aldosterone and ecdysone to induce *prc* mRNA expression, elevate proteinuria and inhibit nephrocyte filtration (Fig 3B, D, F). Likewise, *dopEcR* in cardiomyocytes is required for aldosterone and ecdysone to induce excess pericardin protein production. Whereas, both hormones were able to stimulate pericardin production in flies with nephrocyte-specific *dopEcR* knockdown (*sns*-Gal4>*dopEcR*(RNAi)), or in flies with nephrocyte-specific and cardiac-specific *EcR* knockdown (*sns*-Gal4>*EcR(*RNAi) and *tin*Δ4-Gal4>*EcR*(RNAi)) (Fig 3I, J, K).

**Figure 3.**
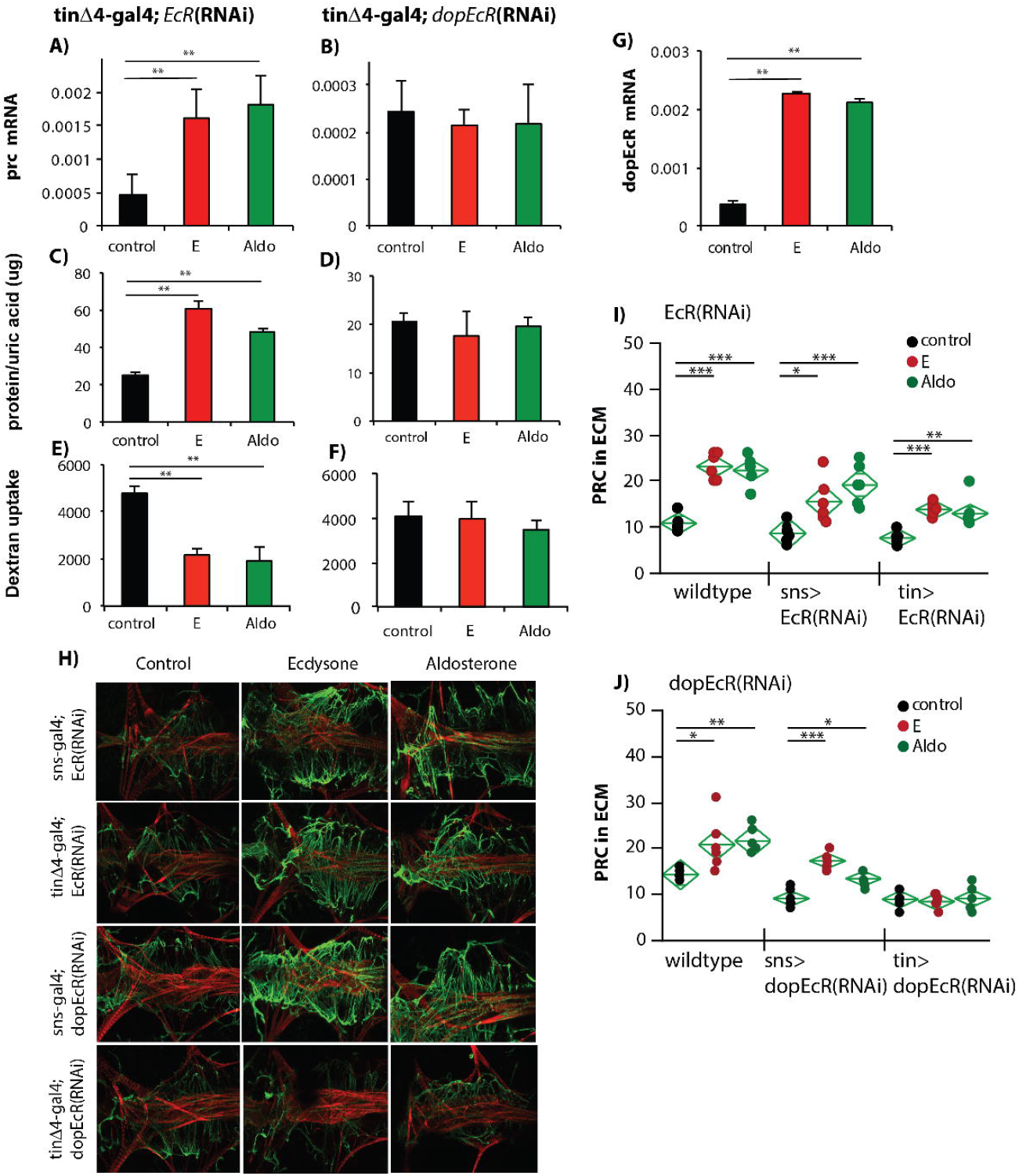
Cardiomyocyte *dopEcR* required for steroid induction of fibrosis and renal pathology. **A, C, E)** Depletion of nuclear hormone receptor EcR by RNAi does not block ecdysone and aldosterone induction of nephrocyte-cardiac tissue *prc* mRNA (A), proteinuria (C) or loss of nephrocyte dextran filtration (E). **B, D, F)** Depletion of GPCR *dopEcR* by RNAi blocks ecdysone and aldosterone induction of nephrocyte-cardiac tissue *prc* mRNA (B), proteinuria (D) and loss of nephrocyte dextran filtration (F). **G)** dopEcR mRNA occurs and is elevated in nephrocyte-heart pericardial tissue of 3 w old adults treated overnight with ecdysone or aldosterone. **H)** Confocal images (representative z-stacks) of nephrocyte-cardiac tissue of 3 w old females after two-week treatment as control, or with ecdysone or aldosterone applied to genotypes that deplete *EcR* or *dopEcR* mRNA in nephrocytes (sns-gal4) or cardiomyocytes (*tin*Δ4-gal4). Cardiomyocyte actin stained by phalloiden, red. Pericardin protein of extra-cellular matrix about nephrocytes and heart, green. **I, J)** Quantification of straining intensity for pericardin ECM protein (each group, six independent nephrocyte-heart preparations) for genotypes to deplete *EcR* mRNA (I) or *dopEcR* mRNA (J) in nephrocytes (*sns*-gal4) or cardiomyocytes (tinΔ4-gal4). (A-G, I, J: One-way ANOVA with Dunnett’s comparison relative to control, * p < 0.05, p < 0.01. Plots show mean with SD.)

## Discussion

Derived from cholesterol, mammalian aldosterone is synthesized in the adrenal cortex into a 21-carbon, C21-hydroxyl steroid and controls plasma Na+ and K+, water balance and blood pressure. In insects, ecdysone has a similar structure - a 27-carbon steroid with hydroxyl groups at C21 and C27. In larvae ecdysone is secreted from the prothoracic gland and controls insect development through its action as 20E via the nuclear hormone receptor EcR. Our data show that both aldosterone and ecdysone fed to adult Drosophila stimulate the expression and accumulation of pericardin in cardiac associated extracellular matrix, and to produce renal dysfunction akin to human fibrosis. In these adults, aldosterone and ecdysone appear to function through the G-protein coupled receptor dopEcR, and not through the canonical nuclear hormone receptor EcR, the otherwise typical pathway by which ecdysteroids regulate insect development and physiology. How aldosterone mimics ecdysone in this context remains unknown. Future work will be needed to determine if aldosterone itself binds to dopEcR, as was shown for Ponasterone A and ecdysone (36), or whether it simply acts as a precursor molecule that can be converted to ecdysone within *Drosophila*. It also remains an open question to determine what roles ecdysone normally plays in development through its control of pericardin, for instance as it might affect heart remodeling during molts and pupation (37).

Although ecdysone is synthesized in the prothoracic gland during development and in adult female follicles, adult somatic tissue including Malpighian tubules also produce ecdysone in response to desiccation stress (38). Ecdysone circulating in adult hemolymph may act at many sites aside from its classic targets of fat body and ovary (39). Notably, dopEcR functions in the fly brain as an alternative ecdysone receptor (34, 35). Here we document *dopEcR* is required in cardiomyocytes to modulate steroid-induced fibrosis. Fibrosis in humans arises from myofibroblasts that secrete extracellular matrix proteins including fibronectins, elastins and collagens (40–42). In *Drosophila*, we show that pericardin appears to be expressed by cardiomyocytes, suggesting these cells have an analogous function to mammalian myofibroblasts. We also show that chronic induction of pericardin by steroid hormones acting at cardiomyocytes stimulates excess ECM accumulation, proteinuria and nephrocyte filtration defects.

DopEcR is a dual agonist receptor (43). In neurons, DopEcR transduces signals from both dopamine and ecdysone to regulate mating behavior and ethanol sensitivity (34, 35). Activation by dopamine induces cAMP-mediated signal transduction. Ecdysone has greater affinity to dopEcR and through unknown mechanisms will displace dopamine and induce alternative signal transduction mediated by MAP kinases (36). Reports are mixed on whether ecdysone also affects cAMP via DopEcR because dopamine alone increased cAMP in Sf9 cells expressing dopEcR (35, 36). In mammalian cells, cAMP can induce PKA to phosphorylate CREB, which then localizes to promoters. Human CREB targets include several collagen genes associated with extracellular matrix (44, 45), and cAMP stimulation suppresses collagen-I expression in a CREB dependent manner (46). Accordingly, we hypothesize that dopamine-cAMP-associated transduction in response to dopEcR may negatively regulate pericardin. If so, dopamine in the absence of steroid hormones may suppress ECM accumulation.

In contrast, ecdysone stimulated DopEcR can signal through dEGFR to ERK1/2 as seen in transfected Sf9 cells and in a neuronal analysis of ethanol induced sedation (34, 36). In mammals, EGFR signaling is broadly implicated in renal fibrosis (47), and these effects may be modulated in part by GPCR crosstalk (48). MAPK/ERK modulates TGF-β1 and its transcription factor Smad2/3, which potently induces collagen transcription in fibrosis (49). Future work can resolve whether aldosterone and ecdysone in Drosophila uses dopEcR to license the ability of dEGFR to stimulate pericardin.

Studies in mammals suggest aldosterone may also signal via a membrane associated GPCR. GPER1 has been proposed to function as a non-genomic aldosterone receptor (21, 22, 50). GPER1-dependent effects induced by aldosterone are reported in various models including renal cortical adenocarcinoma cells (17), and are inferred from mouse models with tissue specific mineralocorticoid receptor gene deletion (51). However, no data establish the mechanism of non-genomic action for aldosterone through GPER1 (23, 52). Furthermore, the leading steroid candidate for GPER1 is 17β-estradiol (53), and the GPER-dependent impact of aldosterone in cells could reflect heterologous desensitization. Alternative dopEcR homologs can be identified in the human genome using the DIOPT Ortholog Prediction Tool, including GRP52 (sequence similarity 46%) and UTS2R (sequence similarity 44%). GPR52 is an orphan G-protein coupled receptor described to modulate Huntingtin protein (HTT) through cAMP-dependent mechanisms (54). Knockdown of Gpr52 reduces HTT levels in a human tissue model, whereas neurodegeneration is suppressed by knockdown of *dopEcR* in transgenic Drosophila that express human *Htt*. The Urotensin II receptor (UTS2R) is a conserved GPCR implicated to function in renal fibrosis by trans-modulating EGFR and activating MAPK (55, 56). In an induced model of rat diabetes, kidneys expressed elevated Urotensin II, and UTS2R was required for exogenous Urotensin to induce TGF-β1 and ECM collagen. These candidates illustrate the translational potential of the Drosophila steroid-induced model of fibrosis. Understanding how dopEcR modulates fibrosis in Drosophila will uncover how related signaling elements affect fibrosis in humans.

## Material and methods

### Fly stocks

Unless noted, wildtype flies were *yw* (ywR). *Tin*Δ4-Gal4 was a gift from the Manfred Frasch laboratory (57). *sns*-Gal4 was obtained from the Bloomington Stock Center (Stock #76160) and UAS-*pericardin*(RNAi) was obtained from the Vienna Drosophila Research Center (Stock #GD <u>41321</u>). UAS-*EcR*(RNAi) was from the laboratory of Neal Silverman (UMass Medical). *dopEcR*(RNAi) is kk103494 of VDRC.

### Steroid and diet treatment

Ecdysone (Sigma-Aldrich #E9004), 20-Hydroxyecdysone (Sigma-Aldrich #H5142) and Aldosterone (Sigma-Aldrich #A9477) were dissolved in ethanol at 5mg/ml. Flies were reared in bottles with emerging adults permitted to mate for 2-3. Adult were then separated by sex into 1L demography cages at ~ 120 adults per cage. Adults were fed standard laboratory cornmeal-yeast-sugar diet until age 7-10 days, at which time food media was switched to 0.5g Genesee Scientific instant fly media (Genesee Scientific #66-117) hydrated with 2ml of water containing vehicle control (150 ul ethanol) or vehicle with 150 ul of hormone solution. For chronic exposure to steroids, flies were treated for the next 14 days at 25 °C fly with media vials changed every 3 days. For overnight exposure to steroids, flies were maintained in demography cages with untreated instant fly media until age 20 days old, then exposed to diets with appropriate hormone conditions for 24 hours. In all trails, renal traits and *prc* mRNA were assessed in adults at 3 weeks old. The same protocols were used to expose adults to high salt or high sugar, where instant media was moistened with water containing 1.5% NaCl. To vary dietary glucose, adults were aged to 3 weeks on otherwise standard lab diet where glucose was set at 5% (control, normal) or at 34% (high sugar diet).

### Proteinuria

For each biological replicate, frass of 15-20 males was collected for 2.5 hours in a 1.5ml centrifuge tube covered with a breathable foam plug, at 25 °C. Deposited frass was fully dissolved with 20ul 1xPBS, providing 10ul to assess total protein and 10ul to measure uric acid, which serves as a proxy for the quantity of deposited frass. Total urine protein was determined by Pierce BCA Protein assay (Thermo Scientific #23227). Uric acid was measured by QuantiChrom Uric Acid Assay (Bioassay systems, DIUA-250).

### Immunohistology

Nephrocyte-heart tissue from 3 w old adults were dissected in PBS, fixed with 4% formaldehyde in PBS for 30 minutes and washed three times for 10 minutes with PBTA (1xPBS,1.5% BSA, 0.3% Tween20) at room temperature. The washed tissue was incubated with 100 ul primary antibody (mouse anti-Pericardin 1:100, Developmental Studies Hybridoma Bank) diluted in PBTA overnight at 4C, washed 3×10 minutes with 1ml PBTA at room temperature, then incubated in secondary antibody (goat anti-mouse Alexa488 1:200, Alexa555-phalloidin 1:100, ThermoFisher Scientific) diluted in PBTA overnight, washed 3×10 minutes with 1ml PBTA at room temperature, and mounted. Confocal images were obtained with a Zeiss 800 and quantified by imageJ software. The full length of pericardial tissue was imaged from all samples at 488 nm with the same laser intensity setting to produce a Z-stack comprised of 46 optical slices.

### Nephrocyte filtration

Adult nephrocyte-heart tissue was dissected in ADH (Artificial Drosophila Hemolymph, 108 mM Na^+^, 5 mM K^+^, 2 mM Ca^2+^, 8 mM MgCl_2_, 1 mM NaH_2_PO_4_, 4 mM NaHCO_3_, 10 mM sucrose, 5 mM trehalose, 5 mM Hepes, pH 7.1), incubated at 25°C for 15 minutes with AlexaFluor568-Dextran (10,000 MW, Life Technology) diluted in ADH at a concentration of 33mg/ml, washed 3×10 minutes with cold PBS at 4C, then fixed in 4% formaldehyde for 10 minutes at room temperature, washed 3×10 minutes with PBS at room temperature, and mounted in PBS. Confocal images were obtained with a Zeiss 800 and quantified by imageJ software.

## Supporting information

Zheng Supplemental data

## Acknowledgements

WZ and MT received support from NIH grant PO1 AG033561, NIDDK Diabetic Complications Consortium grant DK076169, and the office of the Dean of Biology and Medicine, Warren Alpert School of Medicine, Brown University. KO received support from NIH grant R01 HL132241. Erika Taylor assisted KO at the SBP Medical Discovery Institute.

## Supplemental materials

**Fig S1.** 24 h aldosterone treatment does not induce proteinuria.

**Fig S2.** Validation of RNAi knock-down (*prc, dopEcR, EcR*).

**Table S1.** Primers

