## Supplementary material for "Extra-cellular matrix induced by steroids through a G-protein coupled receptor in a Drosophila model of renal fibrosis": Zheng Supplemental data

Figure S1

Proteinuria is not induced by feeding aldosterone to 3 week old males. (Nor when aldosterone is fed overnight to young males, data not shown.)

Proteinuria from overnight aldosterone treatment

|  | protein/uric acid (ug) |
| --- | --- |
| control | 26.7417 |
| aldo | 35.1813 |

Fig S3

**Validation of RNAi efficiency.** Means, with standard deviation, of qPCR (normalized relative to rp49) from full nephrocyte-myocardial tissue, three replicates genotype, dissected from females at 20 days old. Differences between wildtype and RNAi are significant at least at p < 0.02 in each case.

tinD4>prc(RNAi)

|  | prc mRNA |
| --- | --- |
| wildtype | 0.000267956 |
| prc(RNAi) | 0.000148813 |

tinD4>dopEcR(RNAi)

|  | dopEcR mRNA |
| --- | --- |
| wildtype | 0.000383962 |
| dopEcR(RNAi) | 9.0084e-05 |

tinD4>EcR(RNAi)

|  | EcR mRNA |
| --- | --- |
| wildtype | 0.000700363 |
| EcR(RNAi) | 0.000240218 |

Table 1. Primers for qPCR.

**Rp49**

F: 5’- GCA CTC TCT GTT GTC GAT ACC CTT G -3’

R: 5’- AGC GCA CCA AGC ACT TCA TC -3’

**Pericardin**

F: 5’- CGG AGG ACA GGC TAC AAT AAG -3’

R: 5’- TTC CAG GCT GAG TTT CGT ATC -3’

**Col4a1**

F: 5’- GCT CTG TGC GAT TTG AGT TTG -3’

R: 5’- CTT CTG CTC CCT TGA ATC CTT -3’

**Viking**

F: 5’- GAT CTA CGA CAA CAC TGG TGA G -3’

R: 5’- TTC GCC ACG AAG TCC AAT AG -3’

**EcR**
F: 5'-TGA AGA CTC CTA TGC TGC-3'

R:5'-CGA CGT TGT GCT TCG TAA-3'

**dopEcR**

F: 5’- CTT AGG TCC CAG CCT CAT TTC -3’

R: 5’- AGC CAG AGC AGT TGC ATA TT -3’
